# Advancing EDGE Zones: spatial priorities for the conservation of tetrapod evolutionary history

**DOI:** 10.1101/2023.08.15.553233

**Authors:** Sebastian Pipins, Jonathan E.M. Baillie, Alex Bowmer, Laura J. Pollock, Nisha Owen, Rikki Gumbs

**Affiliations:** On the Edge, London, UK; Royal Botanic Gardens, Kew, London, UK; Department of Life Sciences, Imperial College London, Ascot, Berkshire, UK; Science and Solutions for a Changing Planet DTP, Grantham Institute, Imperial College London, London, UK; Department of Global Health & Development, London School of Hygiene and Tropical Medicine, London, UK; Department of Biology, McGill University, Montreal, Quebec, Canada; Quebec Centre for Biodiversity Sciences, Montreal, Quebec, Canada; EDGE of Existence Programme, Zoological Society of London, London, UK

## Abstract

The biodiversity crisis is set to prune the Tree of Life in a way that threatens billions of years of evolutionary history. To secure this evolutionary heritage along with the benefits it provides to humanity, there is a need to understand where in space the greatest losses are predicted to occur. We therefore present threatened evolutionary history mapped for all tetrapod groups, globally and within Biodiversity Hotspots, and identify priority regions of Evolutionarily Distinct and Globally Endangered (EDGE) species at both a grid cell and national level. We find that threatened evolutionary history peaks in Cameroon, whilst EDGE species richness peaks in Madagascar. We refined and advanced the 2013 EDGE Zone concept for spatially prioritising phylogenetic diversity using a novel complementarity procedure with uncertainty incorporated for 33,628 tetrapod species. This involved using extinction risk, phylogenetic, and spatial data to iteratively select areas with the highest accumulated threatened evolutionary history driven by unique species compositions. We identify 25 priority EDGE Zones, which are insufficiently protected and disproportionately exposed to high levels of human pressure. Together, the 25 EDGE Zones occupy 0.723% of the world’s surface but harbour one-third of the world’s threatened evolutionary history, half of which is endemic to these grid cells. They also contain part of the distribution of 918 EDGE tetrapod species, representing near one-third of all EDGE species, with 480 being endemic. Our tetrapod EDGE Zones highlight areas of immediate concern for researchers, practitioners, policymakers, and communicators looking to safeguard the Tree of Life.

## Introduction

With over half of Earth’s land surface exposed to high levels of human pressure (1) and the current protected area network not sufficient to avert the present extinction crisis (2, 3), determining which areas should be prioritised for conservation is key. Typically, prioritisations have reflected patterns of endemism, extinction risk, and species richness (4, 5), and tend to overlook spatial patterns of evolutionary history, an important (6–8) and largely neglected (9–11) component of biodiversity. Evolutionary history is being disproportionately lost in a way that is threatening deep evolutionary branches across the Tree of Life (12, 13). This entails not only a loss in our natural heritage but also a loss in the diversity of biological features and functions resulting from divergent evolutionary histories (14). Sets of taxa containing a greater variety of evolutionary histories provide a greater variety of benefits to humankind (15, 16); highly evolutionarily distinct species in particular often show multiple benefits for humanity, providing numerous services such as food, medicine, and spiritual connections (17). Maintaining evolutionary history also preserves living variation, providing ‘option value’ in the form of unexplored or potential benefits for future generations (18–20). Identifying and conserving areas harbouring large accumulations of threatened evolutionary history is therefore essential for sustaining natural diversity and human well-being.

The dynamics of evolutionary history have been mapped in several ways, based on the measure Phylogenetic Diversity (hereafter, PD) which sums the lengths of phylogenetic branches to quantify the evolutionary history within and between species (18). Whilst spatial patterns of PD are intrinsically correlated with species richness (21–23), the two aspects of biodiversity can decouple locally (15, 24) and highlight non-congruent priority areas (25, 26). Hotspots of range-restricted PD, referred to as ‘phylogenetic endemism’ (27), display low levels of protection (28) and tend to occur in regions of high human pressure (28, 29). This pattern of low protection is surprisingly common amongst important areas of evolutionary history (26, 30) and, in the Americas, the protected area network captures less terrestrial vertebrate PD than expected by chance (31). In contrast, Biodiversity Hotspots (4) may serve as better surrogates for phylogenetic diversity (31, 32)). Biodiversity Hotspots were not designed with PD explicitly in mind, however, and no quantification of their varying evolutionary histories for all tetrapod groups (i.e., amphibians, birds, mammals, and reptiles) has been carried out.

Spatial priorities of evolutionary history can also be revealed by taking a species-specific approach and identifying areas containing high concentrations of the threatened species who contribute disproportionately to the Tree of Life. One way of valuing species in this way is through the EDGE metric, which ranks species based on a combination of their Evolutionary Distinctiveness (ED) and their Global Endangerment (GE) (33). These ranked lists exist for a variety of taxonomic groups, including mammals (33, 34), amphibians (35), birds (36), corals (37), reptiles (38), gymnosperms (39), and sharks and rays (40). The EDGE approach has been updated to incorporate phylogenetic complementarity (41), which describes how the irreplaceability of focal species is influenced by the extinction risk of closely-related species (42, 43). Species with many secure close relatives are considered less irreplaceable than those with few and highly threatened relatives. As such, EDGE scores now quantify threatened evolutionary history, with the scores representing the amount of PD expected to be lost, in millions of years (MY), that can be averted with conservation action (41). Whilst all species have an EDGE score, only those that: 1) are in an IUCN threatened category (Vulnerable, Endangered, Critically Endangered) or are classified Extinct in the Wild; and 2) have an EDGE score above the median for all species in their clade with >95% certainty, are considered priority EDGE species (41).

There have been several studies mapping the distributions of EDGE species (10, 23, 30, 44), although few consider a wide scope of taxonomic groups (but see (30)). Previously, EDGE species have been used to identify priority regions of mammal and amphibian evolutionary history, termed ‘EDGE Zones’ (23). In our study, we also produced a set of priority sites, focussing on high accumulated levels of unique threatened evolutionary history using a refined methodology that accounts for uncertainty in the data and that now incorporates all tetrapod groups. We consider our approach an update to and extension of Safi et al.’s prioritisation, with the same principal aim of conserving evolutionary history. As such, we retain the term ‘EDGE Zones’ for the priority regions identified.

We also explored patterns of EDGE species richness for all tetrapod groups globally, assessing patterns of endemism and ranking nations by their pooled EDGE richness. We then described patterns of threatened evolutionary history globally and within Biodiversity Hotspots, revealing areas harbouring disproportionate levels of expected loss. Within each EDGE Zone, we quantified the patterns of threatened PD, levels of protection, and extent of human pressure. In formulating these areas, our intention is to highlight global priorities for threatened evolutionary history as an important component of conservation prioritisation, especially in those EDGE Zones currently overlooked by existing conservation efforts, in order to safeguard the Tree of Life.

## Results

### EDGE species richness

We mapped the distributions of 2,937 EDGE tetrapod species (919 amphibians, 683 birds, 618 mammals, and 717 reptiles), representing 98.2% of tetrapods that meet the EDGE species criteria under the novel EDGE2 methodology (2,992 spp. total) (41). Their collective distribution covers 92.9% of the world’s terrestrial surface (Fig 1a). EDGE species richness increases towards the equator (S1 Fig) and is particularly high across large parts of Southeast Asia and the Indo-Gangetic plain, as well as in Hispaniola, the highlands of Cameroon, and the Eastern Arc mountains of East Africa (Fig 1a). Maximum EDGE species richness occurs in Northern Madagascar, with a single 96.5 km × 96.5 km grid cell containing 45 EDGE species, but EDGE species are absent from parts of Central and Southern Australia, Central Europe, and the Sahara Desert.

**Fig 1:**
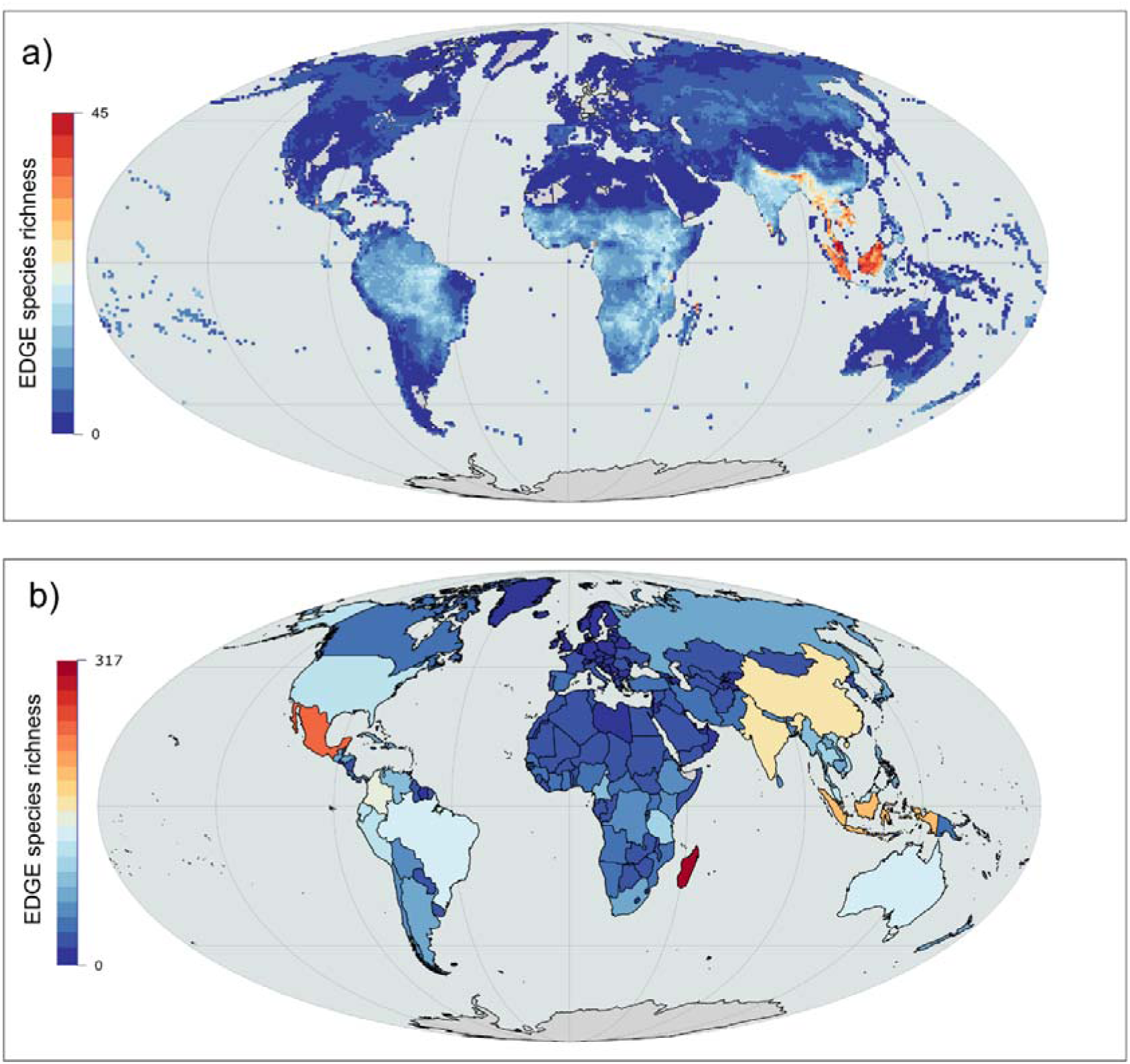
The species richness of Evolutionarily Distinct and Globally Endangered (EDGE) tetrapods. The richness of 2,937 tetrapod EDGE species mapped (a) using a 96.5 km × 96.5 km equal area grid and (b) at the national level in a Mollweide projection.

Madagascar (317 spp.), Mexico (241), and Indonesia (191) contained the highest number of EDGE species (Fig 1b; S1 Data), and 75.6% of EDGE species are endemic to a single country (2,262 of 2,992). However, the pattern varies by taxonomic group; 90.1% of EDGE amphibians are country endemics, followed by 82.4% of EDGE reptiles, 65.5% of EDGE mammals, and 58.5% of EDGE birds. EDGE species endemism is also notable at a grid cell level, with the distribution of 1,451 EDGE species (49.4% of total) limited to any one 96.5 km × 96.5 km grid cell. The patterns also vary for each individual taxonomic group (S2 Fig), although certain areas show co-occurrence for each group (S3 Fig). These include Mesoamerica, the Caribbean, the Andes, the Guinean Forests of West Africa, the Eastern Arc, Madagascar, the Western Ghats, Sri Lanka, and Southeast Asia.

### Threatened evolutionary history

To estimate the amount of threatened evolutionary history that could be secured with conservation action we mapped the summed EDGE scores of 33,628 tetrapod species, representing 92.4% of all 36,376 tetrapod species with EDGE data (Fig 2a; S4 Fig). Several approaches to mapping threatened evolutionary history have involved summing branch lengths from a phylogeny rather than summing species-specific values such as EDGE scores (25, 27–29, 45) (but see (23, 30, 46)). We found an extremely strong positive correlation between both the phylogenetic branch length approach and the summed EDGE score approach for calculating threatened evolutionary history for each tetrapod group (Pearson’s correlation adjusted for spatial autocorrelation (29, 47–49); all ρ > 0.99) (S1 Text), as well as a high overlap in the location of priority grid cells at the 90^th^ (>94.8% overlap) and 95^th^ percentile (>94.4% overlap) for each taxonomic group (S1 Text).

**Fig 2:**
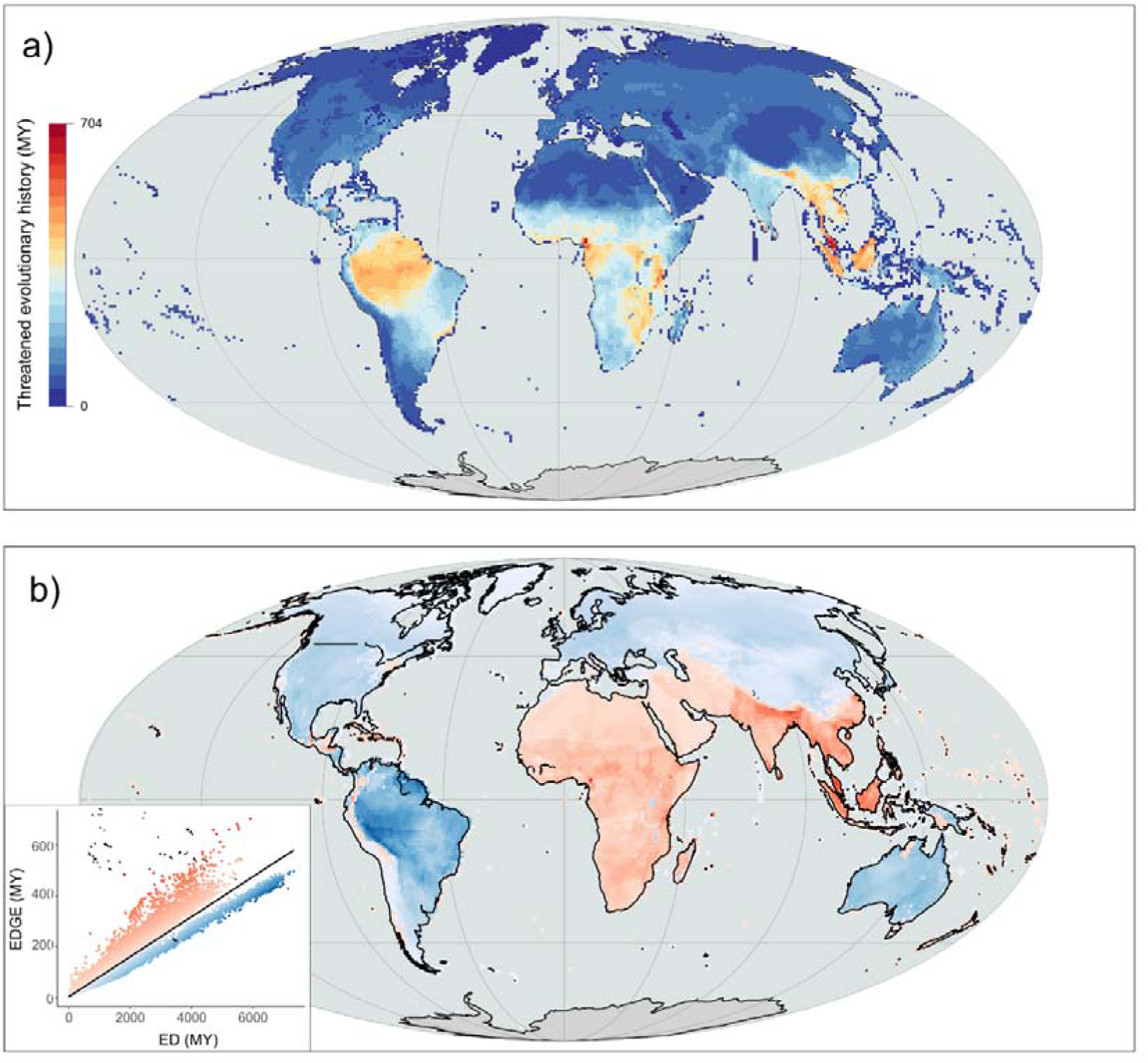
The global distribution of threatened tetrapod evolutionary history. (a) Threatened tetrapod evolutionary history mapped using summed Evolutionarily Distinct and Globally Endangered (EDGE) scores; (b) a linear regression of global patterns of summed EDGE scores against summed Evolutionary Distinctiveness (ED) scores, where warm coloured residuals reflect areas where evolutionary history is more threatened than expected given the summed ED of the region, and cool coloured residuals reflect where evolutionary history is less threatened than expected given the summed ED.

As with EDGE species richness, threatened evolutionary history increases towards the equator (S1 Fig). Southeast Asia, Africa and the Indian subcontinent have a higher incidence of extinction risk relative to the evolutionary history present compared with areas such as South America, where the proportion of evolutionary history that is threatened is lower (Fig 2b). The top-scoring grid cell is in Cameroon, with 703 MY of threatened evolutionary history. Other areas of importance include Honduras, Hispaniola, the Atlantic Forest, the Western Ghats of India, and Sri Lanka.

The distribution of threatened evolutionary history was highly correlated with species richness (Pearson’s correlation: r = 0.916, e.d.f. = 36.3, p < 0.0001). However, priority regions resulting from the two measures are more dissimilar than the high correlation suggests, with the congruence decreasing at high percentiles. For example, for the 80^th^ percentile (i.e., the 20% of highest scoring grid cells), the two have an 84.8% overlap. However, at the 90^th^ percentile this decreases to an overlap of 69.8%, followed by 59.5% at the 95^th^ percentile and 49.4% at the 97.5^th^ percentile. Areas of high species richness dominate large parts of the Amazon basin and the Atlantic Forest, reflecting widely distributed shared species compositions (S5 Fig). Meanwhile, areas of high threatened evolutionary history at the 97.5^th^ percentile show a wider array of geographic locations, including Southeast Asia, Cameroon, Madagascar, the Eastern Arc, the Western Ghats, and Sri Lanka (S5 Fig). This dissimilarity in high scoring areas underlines the importance of considering multiple facets of biodiversity within conservation prioritisations (30, 50, 51).

### EDGE Zones

To identify a complementary set of grid cells that capture irreplaceable and threatened evolutionary history, we iteratively selected grid cells based on their total threatened evolutionary history, removing the species found in each grid cell prior to the next iteration (see methods). We captured 32 priority grid cells across which the constituent species captured 25% of total threatened evolutionary history (our chosen threshold). The complementarity procedure was then repeated 1000 times using a distribution of EDGE scores to account for the uncertainty in phylogenetic, extinction risk, and spatial data. The frequency of which a grid cell is selected in this way gives an indication of how confident we are, given data inadequacies, that this site contains a significant amount of threatened evolutionary history not found elsewhere (termed irreplaceability, measured between 0 and 1). The irreplaceability of the initial 32 priority cells had a median value of 0.79 (S6 Fig); four grid cells, found in Northern Madagascar, Hispaniola, Mexico, and Sri Lanka, were selected in every one of the 1000 iterations and a further nine grid cells were selected in more than 90% of iterations. The uncertainty analysis altogether selected from a pool of 320 grid cells, ranging from an irreplaceability of 0.001 to 1 and with a median of 0.007 (S6 Fig).

Geographically proximate clusters of priority cells were then grouped and joined with contiguous cells (those neighbouring from any direction) from the uncertainty analysis (S6 Fig). Eight clusters, in Sri Lanka, Seychelles, Southern Madagascar, Central Madagascar, Ethiopia, Colombia, Mexico, and the Atlantic Forest of Brazil, were unaffected by this and did not increase in size. The remaining 17 clusters were extended by the inclusion of contiguous cells, by up to a maximum of 14 additional cells in the case of the Gangetic Plains. Across the priority clusters, we found that both the proportion of endemic species and the summed EDGE score were significant predictors of the irreplaceability values of the grid cells within (S2 Text). Overall, this resulted in 25 ‘EDGE Zones’ comprising 112 grid cells (Fig 3). These EDGE Zones are spread across five continents and 33 countries, covering 0.723% of the Earth’s terrestrial surface. They are located over 117 ecoregions and 10 different biomes, with 23 EDGE Zones overlapping with the Tropical & Subtropical Moist Broadleaf Forests biome, eleven overlapping with Mangroves, and eight overlapping with Tropical & Subtropical Dry Broadleaf Forest. Of the 109 developing countries with global Multidimensional Poverty Index data (52), which assess a person’s combined deprivation in health, living standards, and education, 29 of these overlap with EDGE Zones. The score for these countries falls between the 6^th^ and 94^th^ percentile, with the median being the 42^nd^ percentile.

**Fig 3.**
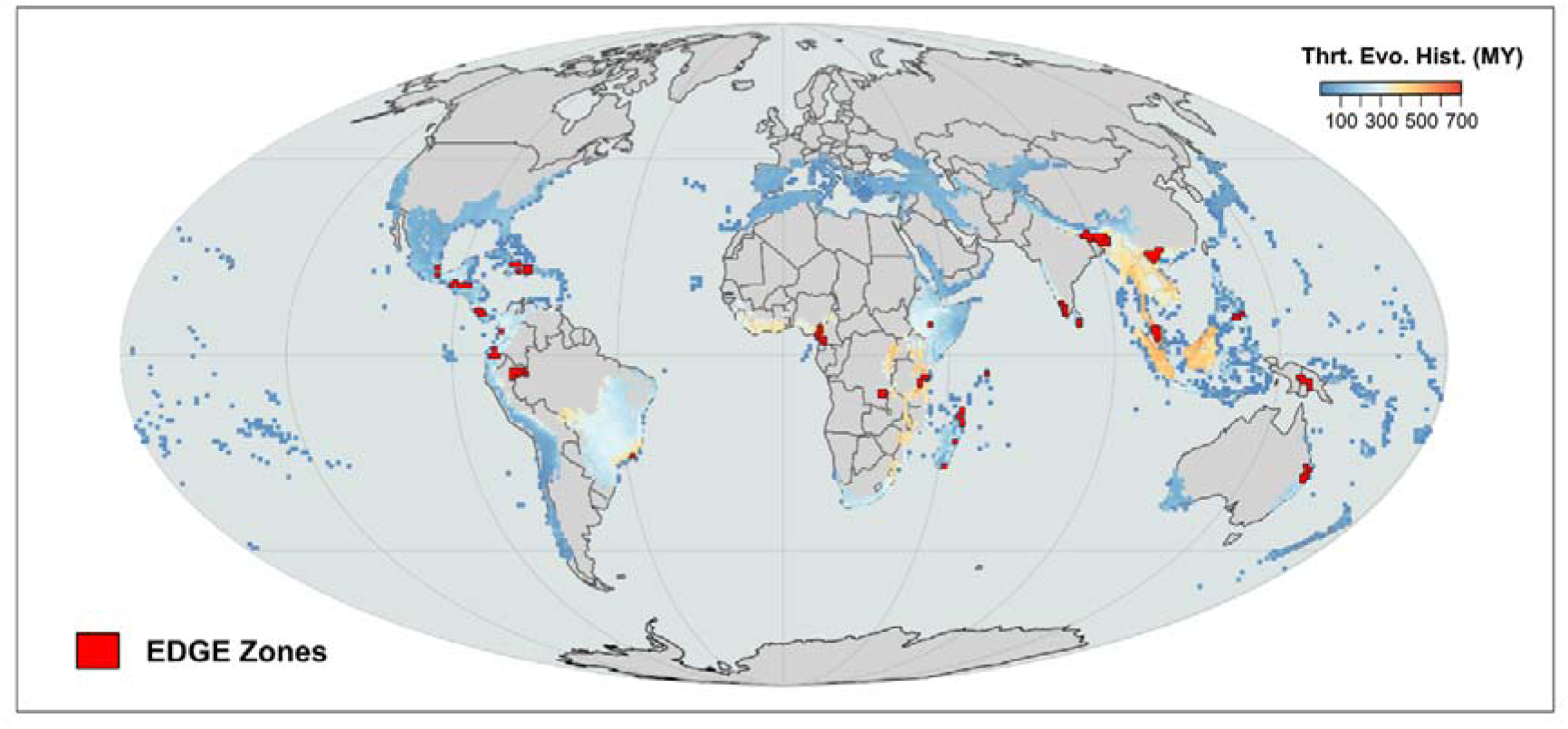
EDGE Zones. The locations of EDGE Zones, a complementary set of 25 grid cell clusters containing large and unique accumulations of threatened evolutionary history. Here, EDGE Zones are superimposed over existing Biodiversity Hotspots, where colours represent per grid cell threatened evolutionary history, increasing from shades of dark blue through to orange.

The priority cells selected in our EDGE Zones analysis (referred to here as EDGE Zone priority cells) were robust to resolution size, mode of calculation, and metric choice: 83% of grid cells selected using a coarser resolution of 193 km × 193 km were overlapping or contiguous to EDGE Zone priority cells (S7 Fig); there was a 97.7% overlap between EDGE Zone priority cells and cells selected using phylogenetic branch-length calculations of expected PD loss (S8 Fig), with the two methods showing a strong correlation in the frequency in which cells were selected (ρ = 0.958, p < 0.0001); there was a 68.8% overlap between EDGE Zone priority cells and cells selected using EDGE scores weighted by range size (‘EDGE rarity’; S9 Fig).

In total, there are 11,662 tetrapod species found in EDGE Zones (34.7% of total), 1,717 threatened species (26.8%), and 918 EDGE species (31.3%) (S1 Table; S2 Data). In every case, this is more than the random expectation; EDGE Zones capture 40% more species, 437% more threatened species, and 348% more EDGE species than would be expected were EDGE Zones distributed at random. The Zone with the most EDGE species is Northern Madagascar with 122 (Fig 4a), 29.8% more than the next highest scoring Zone in Hispaniola with 94.

**Fig 4:**
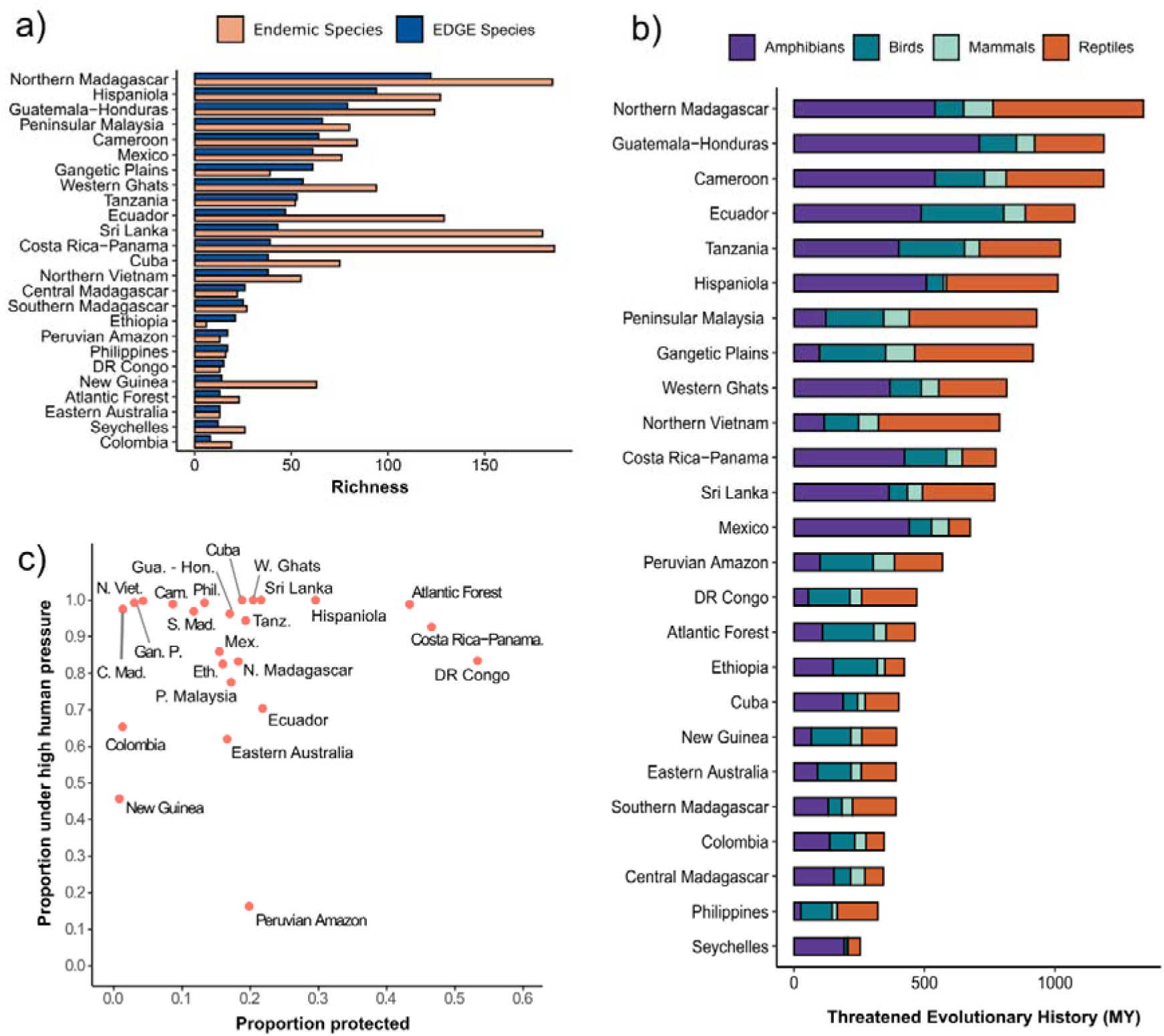
The diversity, human pressure, and threatened evolutionary history within EDGE Zones. (a) The richness of EDGE species and endemics species within EDGE Zones; (b) The threatened evolutionary history (in millions of years; MY) across different EDGE Zones, calculated using the combined EDGE scores of all species found within each Zone and displayed in terms of the different tetrapod groups; (c) the proportion of each EDGE Zone that is under any form of protection and that is experiencing high levels of human pressure. The Seychelles EDGE Zone was excluded from (c) due to absent human footprint data in this location.

Together, if all the species that have at least a part of their range in an EDGE Zone were saved, this would secure 33.3% of total threatened tetrapod evolutionary history (equivalent to 16,820 MY, and 265% more than the 6,345 MY if EDGE Zones were drawn at random). This includes 37.2% of threatened bird evolutionary history, along with 34.3% for amphibians, 31.6% for reptiles, and 31.4% for mammals. Again, Northern Madagascar is the EDGE Zone with the largest accumulation of threatened evolutionary history with 1,339 MY, followed by Guatemala-Honduras with 1,188 MY (Fig 4b).

Naturally, the size of EDGE Zones mean they do not capture the full ranges of all constituent species; for those non-endemic species found within EDGE Zones, the proportion of their ranges contained within the 25 EDGE Zones is a median of 13.5% for amphibians, 6.96% for reptiles, 4.19% for mammals, and 3.57% for birds. Collectively, EDGE Zones contain a median of 4.73% of the ranges of the non-endemic tetrapod species found within. However, endemism levels are notable; EDGE Zones harbour 1,457 grid cell endemics, which is 2,749% more than the random expectation of 53 endemic species. Furthermore, 1,727 species are endemic to anywhere within an EDGE Zone (14% of all species found within an EDGE Zone), meaning 15.5% of total threatened tetrapod evolutionary history is contained exclusively within EDGE Zones. The most endemics are found in Costa Rica-Panama (186), Northern Madagascar (185), and Sri Lanka (180) (Fig 4a). Furthermore, 52.2% of EDGE species in EDGE Zones are endemic to a single Zone (480 of 918 species), equating to 16.0% of all EDGE tetrapods (480 of 2,992).

Using the Human Footprint Index (53), we then explored the proportion of EDGE Zones experiencing appreciable levels of human disturbance (scores of above 4 or more, see methods) (Fig 4c). We found that 81.5% of the total area covered by EDGE Zones is under high human pressure and that this varies by EDGE Zone; from a low of 16.2% to a high of 100%, with the median level of high human pressure in EDGE Zones being 85.3% (S1 Table). Three EDGE Zones (Sri Lanka, Cuba, and the Western Ghats) are entirely under high human pressure. Between 1993 and 2009, a mean shift of 7.23% of the land within EDGE Zones transitioned from low to high human pressure; but this also varied, with 5.8% of the land within Peninsular Malaysia shifting to lower human pressure, compared to 32.8% of the area within New Guinea shifting to high human pressure. The human pressure in EDGE Zones is higher than the background expectation of 56.6%.

A mean of 20% of EDGE Zone land is under any form of protection (*relaxed* protection), whilst 10% is under *stricter* protection standards (IUCN I:IV) (Fig 4c). When assessing the protection standards within each EDGE Zone grid cell, we found levels of relaxed protection were comparable to the United Nations Convention on Biological Diversity’s (CBD) Aichi target of 17% (*t*(111) = −0.232, p = 0.817) (54), but were significantly below the target when based on stricter protection standards (*t*(111) = −6.73, p < 0.0001). Both relaxed and stricter protection levels were significantly below the CBD’s Kunming-Montreal 2030 milestone of 30% (*t*(111) = −8.11, p < 0.0001; *t*(111) = −16.5, p < 0.0001) (55). DR Congo saw the highest cover of any form with protection at 53%; New Guinea and Colombia have less than 1% of their area protected, and four Zones (New Guinea, Colombia, the Western Ghats, and Ecuador) show zero protection when stricter protection categories were considered. However, in some cases these scores are an artifact of inconsistencies in national data reporting, given that India withholds data on 900 protected areas, influencing the low score seen in the Western Ghats (https://www.protectedplanet.net/country/IND). Even within the protected portions of EDGE Zones, high human pressure levels are found across 75.7% of their extent, rising to 83% for the non-protected portions.

Within EDGE Zones, amphibians have significantly greater median EDGE scores per grid cell compared with the other tetrapod classes (p < 0.001 for all pairwise comparisons made using ANOVA with Tukey’s Honest Significant Difference Test; S3 Text), but outside of EDGE Zones, the median EDGE scores of reptiles are significantly greater (P < 0.0001). Relative to their richness within EDGE Zone grid cells, however, there was no difference between the amphibian and reptile proportional contribution to threatened evolutionary history (p = 0.998), although both contributed significantly more than mammals and birds (p < 0.0001; Fig 5; S3 Text).

**Fig 5:**
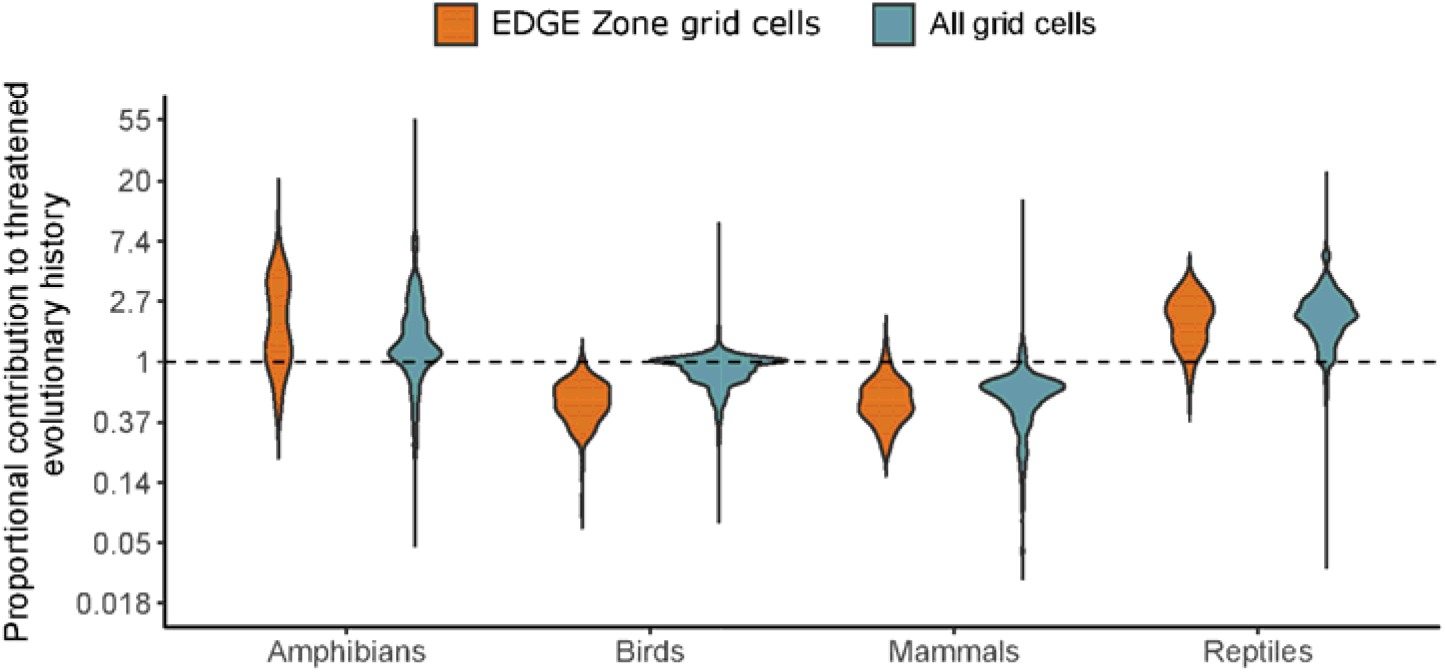
The proportional contribution of tetrapods to the threatened evolutionary history within and outside of EDGE Zone grid cells. The proportional contribution of each tetrapod group to the threatened evolutionary history of EDGE Zone grid cells and all grid cells, where a score of >1 means that a group contributes more to the threatened evolutionary history of a cell than expected from its per cell richness: for example, if amphibians represent 30% of a cell’s species richness but contribute 60% to threatened evolutionary history, their proportional contribution is 2.

### Biodiversity Hotspots

Biodiversity Hotspots, as defined by Myers et al. (4), cover 18.5% of the world’s terrestrial surface, capturing 73.5% of EDGE tetrapods and 74.5% of tetrapod threatened evolutionary history (S10 Fig). The distribution of this diversity is unevenly split across the 36 different Hotspots (S2 Table). For instance, the three Hotspots with the highest median threatened evolutionary history (relating to the median EDGE scores of grid cells found within each hotspot) are Sundaland (393 MY), Indo-Burma (354 MY), and the Guinean Forests of West Africa (339 MY). These same three hotspots scored highest for peak threatened evolutionary history (the maximum summed EDGE grid cell found in each hotspot), with the Guinean Forests of West Africa ranked first (Fig 6). In terms of EDGE tetrapod richness, Madagascar and the Indian Ocean Islands (334 spp.) ranked first, followed by the Caribbean (300 spp.), and Mesoamerica (297 spp.). All three metrics considered were positively correlated with species richness (maximum summed EDGE – r(34) = .6, p = .0001; median summed EDGE – r(34) = .45, p = .0001; EDGE tetrapod species richness – r(34) = .64, p < .0001).

**Fig 6.**
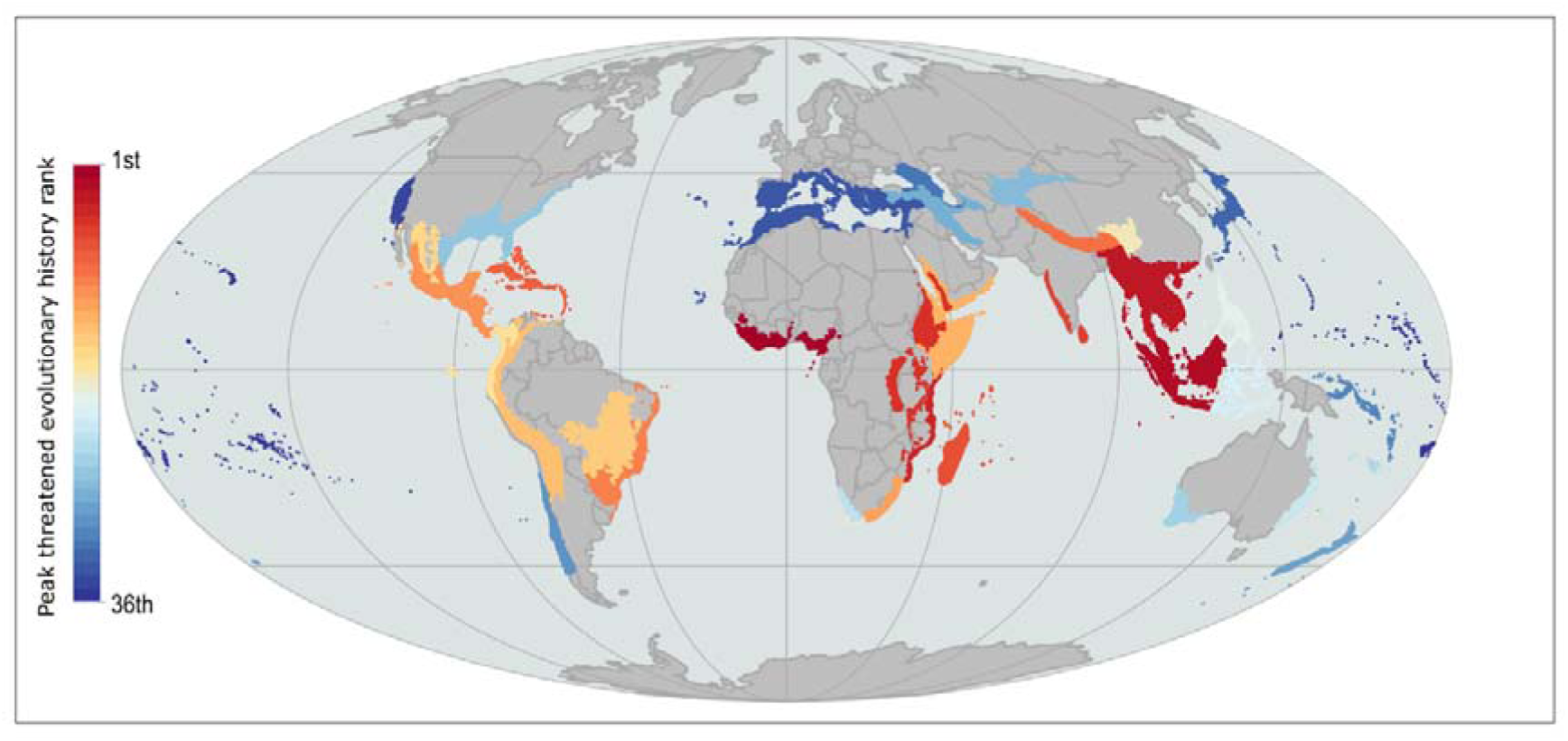
Biodiversity Hotspots ranked by peak threatened evolutionary history. The rank of all 36 Biodiversity Hotspots, as defined by Myers et al. (2000), was measured by comparing the grid cells in each with the largest amounts of threatened evolutionary history and coloured from first in dark red to last in dark blue.

## Discussion

This study is the first to map EDGE species from all tetrapod groups, revealing a high endemicity at both a national and grid cell level. Spatial patterns of threatened evolutionary history diverge from species richness at the highest-ranking priority grid cells, with key areas for threatened evolutionary history including Cameroon, Tanzania, and Hispaniola. To drive the conservation of threatened evolutionary history, we identified 25 priority regions of disparate communities of species, termed ‘EDGE Zones’, which together harbour 33.3% of threatened evolutionary history and 31.3% of EDGE tetrapods, with approximately half of both being found nowhere else. Concerningly, high levels of human pressure are ubiquitous across these Zones, and protection levels largely fail to meet globally recognised targets for 2020 and 2030 (55).

Given Madagascar’s highly endemic biodiversity (56), phylogenetic importance (28, 30), and high levels of extinction risk (57), it is unsurprising that the island harbours the greatest number of priority EDGE tetrapods (10.4% of total). This includes important EDGE species such as the 2^nd^ top ranked EDGE reptile, the Madagascar Big-headed Turtle (*Erymnochelys madagascariensis* ; EDGE score: 84 MY), and the 2^nd^ top ranked mammal, the Aye-aye (*Daubentonia madagascariensis* ; EDGE score: 20 MY). However, high micro-endemism within Madagascar (58, 59) means the EDGE richness here is isolated in patches along the Northern and Eastern parts of the island. In contrast, very large areas of Southeast Asia show elevated levels of EDGE species, reflecting how the ‘looming biodiversity disaster’ in this region is impacting highly unique and wide-ranging species across extensive parts of their range (60).

Half of EDGE tetrapods are endemic to any one grid cell and three quarters are found exclusively in a single country. Given that environmental policy is usually set at the national level (61), our findings suggest that nations must show leadership in protecting their most evolutionarily distinct species. By protecting EDGE species locally, and thus their habitat and other sympatric species, nations can not only preserve their unique biodiversity and the benefits provided by it, but they can also safeguard the global values captured by maintaining the Tree of Life (8, 16).

Whilst threatened evolutionary history shows elevated levels across large swathes of the tropics, it is disproportionately high in parts of Africa, Southern Asia, and Southeast Asia. This reflects highly distinctive and threatened concentrations of tetrapods, which drive greater levels of threatened evolutionary history than would be expected from their species richness alone. In particular, the phylogenetically diverse groups of reptiles and amphibians (29) contribute significantly to the threatened evolutionary history of grid cells, as seen in Cameroon and Peninsular Malaysia, where threatened evolutionary history peaks.

In representing threatened evolutionary history, we elected to use summed EDGE scores. Whilst it has been found that using species as the unit of evolutionary history can lead to spatially efficient prioritisations for PD (36), many studies have mapped spatial patterns using branch lengths from phylogenies (25, 27–29). In our study we found a high degree of similarity between the two approaches, with both methods being highly correlated and producing near identical priority regions. Within this finding lies an opportunity for policy makers or conservation practitioners: to assess patterns of threatened evolutionary history in an area, they can simply sum publicly available EDGE scores for the species found within. This approach is more accessible to non-specialists both in theory and application and can be computed at a fraction of the time of branch-length-dependent methods, providing an avenue to represent evolutionary history more readily within conservation planning activities. Considering species are the typical unit of conservation, using species-specific scores also allows for the inclusion of other species-level biodiversity measures, such as cultural value and functional traits (51), or the incorporation of species from disparate clades into prioritisation exercises (36).

The expansion of our EDGE Zones to incorporate adjacent areas with similar communities of species based on the uncertainty in the phylogenetic and extinction-risk data ensures that these prioritisations are robustly delineated to highlight areas of utmost concern. We can say this with confidence because the prioritisations were robust to the uncertainty in the underlying data, to method choice of calculating threatened evolutionary history, and to grain size used in the complementarity procedure. Using complementarity also ensured that the Zones captured disparate species compositions and unique evolutionary histories; indeed, they overlap with 117 ecoregions and one-third of all tetrapod species.

Although only a fraction of the ranges of non-endemic constituent species are captured within Zones (~5%), 42% of the threatened species and 52% of the EDGE species found within are endemic. As such, roughly 1/6^th^ of global threatened tetrapod evolutionary history is found exclusively in EDGE Zones, and 1/3^rd^ of threatened tetrapod evolutionary is contained partially within EDGE Zones. EDGE Zones are therefore not intended to be the only sites prioritised for protection, but rather they are key regions where the failure to protect the biodiversity within will have major consequences for the Tree of Life and the benefits it bestows. Further research will be needed to assess the conservation opportunities within each EDGE Zone to find where interventions are possible and most likely to have an impact.

The Cameroon EDGE Zone harbours the grid cell with the greatest accumulation of threatened evolutionary history, in part driven by the dramatic declines of evolutionarily distinct amphibians in the region (62, 63). This includes four Critically Endangered EDGE puddle frogs (*Phrynobatrachus* spp.) and eight highly threatened EDGE egg frogs (*Leptodactylodon* spp.). In fact, there is considerable overlap between EDGE Zones and high concentrations of threatened amphibians (64). These areas of overlap include Mesoamerica, the northern Andes, the Atlantic Forest, Cameroon, the Eastern Arc of Tanzania, Madagascar, the Western Ghats, Sri Lanka, and the Philippines. This can be explained by the patterns of high distinctiveness and extinction risk seen in amphibians; relative to their richness, amphibians contribute significantly more to the threatened evolutionary history of EDGE Zones than birds and mammals. The Caribbean, where 84% of amphibians are threatened with extinction (65), is a good example of this association between distinctiveness and imperilment. Here, the selection of the Hispaniola EDGE Zone was driven by amphibians (Fig 4). This includes the influence of the highly speciose (202 spp.), threatened (137 spp.), but also evolutionarily distinctive (median ED = 12 MY) clade of robber frogs (*Eleutherodactylus* spp.), of which there are 42 EDGE species in this one Zone.

EDGE Zones also coincide with other priority areas, with 22 overlapping with Biodiversity Hotspots. Several Biodiversity Hotspots contain multiple EDGE Zones, reflecting the presence of disparate assemblages of threatened evolutionary history within them; this includes Hotspots in Sundaland, Mesoamerica, the Caribbean, and Madagascar and the Indian Ocean Islands. In our comparison of Biodiversity Hotspots, we found that some regions consistently ranked highly in their levels of threatened evolutionary history and EDGE richness. For instance, the Indo-Burma, Sundaland, Eastern Afromontane, Guinean Forests of West Africa, and the Western Ghats and Sri Lanka Biodiversity Hotspots all scored in the top ten for each aspect of evolutionary history we explored. Concerningly, all five of these Hotspots are also ranked in the top third most densely populated Biodiversity Hotspots (>114 people per square kilometre) (66), each has a child malnutrition rate of more than 20% (66), and mounting agroeconomic pressures threaten what little intact vegetation is left in most of these regions (67, 68). This necessitates urgent conservation action, and we believe the comparison provided here can add a useful phylogenetic perspective to this endeavour.

Elsewhere, EDGE Zones coincide with priority locations described in other studies of tetrapod evolutionary history (28, 29, 46). All countries forecasted to experience the greatest losses in evolutionary history due to land-use driven species extinctions (46) contain EDGE Zones. Elsewhere we found that 24 EDGE Zones overlap with hotspots of tetrapod phylogenetic endemism (28), and all 25 overlap with priorities of tetrapod human-impacted phylogenetic endemism at the 95^th^ percentile (29). Thus, although EDGE Zones were not formulated based on patterns of endemism, they also effectively capture range-restricted PD.

Our study echoes previous findings reporting low levels of protection and high human impact across phylogenetically important areas (28–30), with 60% of EDGE Zones being covered by less than 10% of strictly designated protected areas and 76% showing north of 80% of their land under high levels of human pressure (Fig 4). Furthermore, human populations found within EDGE Zone countries face appreciable deprivation in education, health, and living standards, as measured using the multidimensional poverty index. One particularly concerning case study surrounds New Guinea, a region long recognised for its high biodiversity and relatively intact primary vegetation (69). We found that one-third of the available land in this Zone saw a shift from low to high human pressure in a 17-year period, driven by expansive deforestation efforts (70) that are projected to cause major losses in the evolutionary history (46). Whilst the protected area coverage across EDGE Zones currently fails to meet the 2030 target of 30% protection, we did find that relaxed-protection levels were comparable to the Aichi target of 17%. Our research demonstrates that large gains of biodiversity are possible within relatively small additions to the global protected area network. As conservation seeks to protect 30% of Earth’s land by 2030, we emphasise that evolutionary history must be considered; more research is needed to build on this and other work (25, 28, 46) to identify best path forward for these areas.

The downstream utility of global mapping exercises such as that presented here have recently been called into question due to the lack of clear translation to action on the ground (71). We believe this is a valid criticism, given that the realised or potential impacts of global mapping research is not commonly reported in the scientific literature. However, we foresee these priority regions guiding future efforts to save the world’s most distinctive and imperilled tetrapod species through various downstream uses. First, the EDGE Zones presented here will guide the activities of the charitable organisation On the Edge (www.ontheedge.org), directing their conservation grant-making, regional campaigns, and grantee-led storytelling. Second, EDGE Zones will form part of the decision-making for resource allocation for the Zoological Society of London’s EDGE of Existence programme (www.edgeofexistence.org), which has already funded work on over 50 EDGE species found within EDGE Zone countries, with a particular focus on the Gangetic Plains and Cameroon. Third, our method of summing EDGE scores offers the potential for extending this approach to other taxonomic groups such as plants and freshwater fish where data is becoming increasingly available (39, 72–74). Finally, we hope this analysis will inspire in-depth, fine resolution research into the current and future levels of irreplaceability, protection, human pressure, climate change, and conservation potential in each EDGE Zone to catalyse applied conservation action.

## Conclusion

The initial formulation of EDGE Zones was conducted under the aim of establishing ‘a spatial perspective for an otherwise species-centred conservation initiative’ (23) in reference to the EDGE approach of phylogenetically informed species conservation. In revisiting this concept, we have retained Safi et al.’s original aim but significantly extended their approach through a revised prioritisation methodology and the consideration of all tetrapod groups and their threatened evolutionary history. In doing so, we have revealed global patterns in the distribution of highly imperilled PD, highlighting where the greatest potential losses are accumulating, where EDGE species are concentrated, and which Biodiversity Hotspots are especially important in capturing this heritage. In revealing these patterns and prioritisations, we have provided a useful frame of reference for conservationists, policymakers, and scientific communicators seeking to safeguard the Tree of Life.

## Materials and Methods

### Species distribution, extinction-risk, and phylogenetic data

We obtained species distribution data from the IUCN Red List of Threatened Species (Version 2021.1) for terrestrial and freshwater mammals and amphibians (75), from BirdLife International for birds (Version 2020.1; (76)), and from the Global Assessment on Reptile Distributions for reptiles (Version 1.5; (29)). Distribution data were filtered so that only native, breeding, and resident extents of distributions were used, where relevant. Antarctic distributions were excluded. Range extents marked as ‘Presence Uncertain’ and ‘Extinct’ were removed. Range polygons were then rasterised into a grid format using a resolution of 96.5 km × 96.5 km with a Mollweide equal area projection. We used EDGE scores from Gumbs et al. (74), generated using the updated ‘EDGE2’ approach (41), as well as 1,000 extinction-risk-weighted phylogenetic trees used to calculate those EDGE scores (74). Species names from the distribution data and phylogenetic data were matched using the taxonomic databases referred to in the EDGE2 calculation of Gumbs et al. (74). Our study used only those species with both distribution and EDGE data, resulting in the inclusion of 5,614 mammals (89.8% of total), 6,809 amphibians (84.9%), 10,937 birds (99.5%), and 10,268 reptiles (92.4%), together representing approximately 92.4% of tetrapod diversity. All mapping and analyses took place in R version 4.1.0.

### EDGE species richness and threatened evolutionary history

To explore the distributions of EDGE species, we mapped EDGE tetrapods at a grid cell level. We then explored EDGE species richness at the national level, using national reporting data from the IUCN Red List (75). EDGE species designations were taken from Gumbs et al. (74).

In the ‘EDGE2’ approach, EDGE scores are species-specific scores calculated by summing the lengths of branches connecting a species at the tip of the tree to the root in an extinction-risk-weighted phylogeny (41). As these scores are derived from overall estimations of expected PD loss for the entire clade, it stands that summing these EDGE scores should correlate highly with total expected PD loss of the clade (19). We therefore ran a series of sensitivity analyses for each tetrapod group to explore how well summing EDGE scores reflects the typical phylogenetic branch length approach for calculating expected PD loss. To calculate expected PD loss, we worked with the extinction-risk adjusted phylogenetic trees used in Gumbs et al. (74). We worked on a per-grid cell basis, summing the branch lengths from the root node to the tips of the trees subset to only the species found within a given grid cell. This was repeated for a distribution of 1000 phylogenies for each group and the average expected PD loss grid-cell values were then contrasted with the summed EDGE grid-cell values by using a Pearson’s correlation adjusted for spatial autocorrelation (29, 47–49), and then comparing the overlap in priority grid cells using six percentiles (80^th^, 85^th^, 90^th^, 95^th^, 97.5^th^, and 99^th^).

As this comparison revealed a high congruence and strong correlation (S1 Text), we proceeded to use summed EDGE scores for our analyses. Alongside the precedent of using species-specific scores for spatial prioritisations of threatened evolutionary history (23, 30, 46), we believe there is merit in this choice considering that EDGE scores are publicly available and maintained by the EDGE of Existence programme (77). Given that a species’ EDGE score reflects the amount of the PD for which it is responsible (it’s Evolutionary Distinctiveness, given by an ED score) that is expected to be lost, we ran a linear regression of EDGE ~ ED to see where there is more threatened evolutionary history than expected given the amount of evolutionary history present in a grid cell (29). The residuals along this linear regression were mapped, with positive residuals highlighting areas where we are projected to lose more evolutionary history than expected given the modelled relationship.

We also assessed the ED of species with EDGE data but that were missing distribution data. For these species, ED scores were generally skewed towards mid to lower percentiles for each tetrapod group (S11 Fig), with the median ED for these data-deficient species ranging from the 37^th^ percentile in birds to the 44^th^ percentile in reptiles. We therefore predict that the data gaps within tetrapod species distributions will not significantly alter the results presented here.

We then mapped all 36 Biodiversity Hotspots (4, 5, 78) at a 96.5 km × 96.5 km resolution and compared them in terms of their maximum EDGE score, median EDGE score, number of species, number of EDGE species, and number of genera.

### EDGE Zones

To identify priority locations of unique threatened evolutionary history, we iteratively selected top-ranking grid cells with the highest summed EDGE scores using spatial complementarity. With each iteration, the species found within the top-ranked cell from the previous selection were removed from the underlying dataset and the process repeated until a threshold was passed. Our threshold for how many sites to select in this complementarity procedure was the minimum number whose pooled species composition together represented 25% of tetrapod threatened evolutionary history. Here, threatened evolutionary history was measured using the extinction risk-weighted phylogenetic trees found within Gumbs et al. (74). The threshold was met when the subtree connecting all constituent species found within N priority grid cells had a combined expected PD loss equal to 25% of the total for all tetrapod species. Constituent species were classified as those whose distribution overlapped with a priority cell. Our method selected large accumulations of threatened evolutionary whether driven by a small number of highly distinct species or a larger number of less distinctive, but more threatened, species as both represent important potential losses from the Tree of Life.

A complementary set of 32 priority cells was selected from this procedure, out of a total set of 3001 cells to capture 100% of tetrapod evolutionary history (S12 Fig). Contiguous cells (those touching each other in any direction) were grouped together to form single EDGE Zones. Two adjacent pairs of disjunct cells in Haiti and two adjacent pairs of disjunct cells in Mexico were grouped together, respectively, leaving 25 priority clusters. The choice to group these cells was made because we believe their geographical proximity would make it impractical to treat these regions separately from a conservation perspective.

To determine how robust the selection of our priority cells was to changes in resolution size, mode of calculation, and metric choice, we repeated our complementarity procedure using (1) a coarser resolution of 193 km × 193 km, (2) the phylogenetic branch length-based calculation of expected PD loss, and (3) ‘EDGE rarity’, relating to a species’ EDGE score divided by the number of grid cells with which its range overlaps.

The complementarity procedure was repeated 1000 times using the distribution of EDGE scores to account for uncertainty. The irreplaceability (the frequency of selection) of the 32 priority grid cells was recorded. Proximate priority grid cells were grouped with contiguous cells from the uncertainty analysis and the resultant clusters were called EDGE Zones. We then ran a General Linear Model to compare which variables significantly predicted the irreplaceability values within these EDGE Zone grid cells.

We explored the total number of species, EDGE species, threatened species, and endemic species (both grid cell endemics and zone-endemics) found in each EDGE Zone and compared the richness of these groups to a random selection of grid cells of a set size equal to the number of EDGE Zone grid cells (the random expectation). We then quantified the proportion of each species’ total range that is contained within EDGE Zones and described the biome-type and number of intersecting ecoregions (79). Using the global Multidimensional Poverty Index (52), we reported the deprivation faced by people within EDGE Zone countries. Using ANOVA with Tukey’s Honest Significant Difference Test, we also tested which tetrapod groups have the greatest median ED scores, median EDGE scores, and the highest proportional contribution to the threatened evolutionary history in each grid cell relative to their richness, both within and outside of EDGE Zones.

To explore human pressure within each zone, we obtained data on the Human Footprint Index at a 1 km × 1 km resolution for both 1993 and 2009 from (53). The Human Footprint Index is a spatial measure of human pressure on the land, factoring in eight different measures such as the presence of built environments, roads, and crop lands. The combined presence of these measures gives rise to a map of cumulative human pressure, with values ranging from 0 reflecting the lowest possible human pressure to a score of 50 reflecting the highest. We categorised scores of below four as low human pressure and scores of above or equal to four as high human pressure as this threshold is indicative of when species are likely to become threatened by human land-use change (1, 80–82). For each EDGE Zone grid cell, we looked at the proportion of high human pressure scores. We then reported the proportion of each EDGE Zone that had shifted to high human pressure between measurements of the Human Footprint Index (1993 and 2009) and compared the proportion of high human pressure within EDGE Zones to the random expectation.

To explore the level of protection occurring within EDGE Zones, we downloaded data on the protected area (PA) network from the World Database on Protected Area (83). Following WDPA guidelines, PAs whose status was proposed or not reported were removed and, for PAs where only point data was provided, we added a circular buffer zone around each point of a size equal to its reported area. The remaining sites were unionised to dissolve the boundaries between each polygon to prevent the double counting of overlapping areas. We then overlaid the resultant PA polygon with the rasterized grid used in the study and calculated the PA percentage cover of each EDGE Zone. To contrast how the protection status affects the overall coverage, we contrasted strict protection standards using PAs in IUCN categories I-IV (where 1a denotes a strict nature reserve and IV denotes a habitat/species management area) with relaxed standards by using all PAs in the dataset. We then compared the level of protection across all EDGE Zones to two thresholds using t-tests; 17%, reflecting target 11 of the Aichi Biodiversity Targets for 2020 (54), and 30%, reflecting the Kunming-Montreal Global Biodiversity Framework target 3 (55). Finally, we assessed how the human pressure varies between protected and non-protected portions of EDGE Zones.

## Acknowledgements

We thank members of On the Edge, the Zoological Society of London’s EDGE of Existence programme, and the IUCN SSC Phylogenetic Diversity Task Force for their constructive feedback and conceptual input in the making of this research.

## Author contributions

**Conceptualization**: Sebastian Pipins, Jonathan E.M. Baillie, Laura J. Pollock, Nisha Owen, Rikki Gumbs

**Data curation**: Sebastian Pipins, Rikki Gumbs.

**Formal analysis**: Sebastian Pipins, Rikki Gumbs.

**Investigation**: Sebastian Pipins, Rikki Gumbs.

**Methodology**: Sebastian Pipins, Nisha Owen, Rikki Gumbs

**Project administration**: Nisha Owen.

**Supervision**: Nisha Owen, Rikki Gumbs.

**Visualization:** Sebastian Pipins

**Writing – original draft:** Sebastian Pipins.

**Writing – review & editing:** Sebastian Pipins, Jonathan E.M. Baillie, Alex Bowmer, Laura J. Pollock, Nisha Owen, Rikki Gumbs

## Supporting Information

**S1 Data. EDGE species richness by nation.** The number of EDGE tetrapod species found in each nation, based on IUCN national reporting data.

**S2 Data. List of EDGE Species found within EDGE Zones.** The EDGE species found in each EDGE Zone, along with their common names, IUCN Red List IDs, and Red List categories (where LC = Least Concern, NT = Near Threatened, VU = Vulnerable, EN = Endangered, CR = Critically Endangered).

**S3 Data. Raster layer of tetrapod summed EDGE scores.**

**S4 Data. Raster layer of EDGE tetrapod species richness.**

**S5 Data. Shapefile of national-level EDGE tetrapod species richness.**

**S6 Data. Shapefile of EDGE Zones.**

**S1 Code. Prioritisation procedure.** https://github.com/sebpipins/EDGE-Zone-prioritisation/

**S1 Table. The diversity within EDGE Zones.** Here, endemic species refers to the number of species found exclusively within a given EDGE Zone, whilst the number of species found exclusively in any singular grid cell of that EDGE Zone is shown in brackets. The percentage of an EDGE Zone under high human pressure is reported for 2009 and 1993 in brackets. Protection levels represent the percentage of an EDGE Zone that is under any form of protection, with stricter protection standards (IUCN grades I:IV) shown in brackets.

**S2 Table. Biodiversity Hotspots.** Threatened evolutionary history is measured using the summed EDGE scores of grid cells overlapping with Biodiversity Hotspots.

**S1 Fig. Regressions of latitude against threatened evolutionary history and EDGE species richness.** Loess regression (blue) of (a) threatened evolutionary history (given by the median summed EDGE score in millions of years) and (b) the median EDGE richness at different degrees of latitude.

**S2 Fig. EDGE tetrapod species richness.** The species richness of EDGE amphibians, birds, reptiles and mammals.

**S3 Fig. Spatial congruence of EDGE tetrapod groups.** The distribution co-occurrence of EDGE species from four tetrapod groups.

**S4 Fig. Threatened evolutionary history of tetrapod groups.** Threatened evolutionary history is represented in millions of years (MY) and mapped using summed EDGE scores.

**S5 Fig. Comparison of species richness and threatened evolutionary history.** (a) The difference in the standardised scores of threatened evolutionary history and species richness per grid cell, where positive values represent greater standardised threatened evolutionary history (green) and negative values represent greater standardised species richness (brown). (b) The 97.5^th^ percentile of threatened evolutionary history grid cells in red and species richness in blue, with the overlap shown in black. Threatened evolutionary history is measured using summed EDGE scores.

**S6 Fig. The irreplaceability of EDGE Zone grid cells.** The (a) location of the grid cells from the uncertainty analysis, where EDGE Zone priority cells are those identified from the complementarity procedure using median EDGE scores (black). Other highlighted cells are those identified from the complementarity procedure repeated on the distribution of 1000 possible EDGE scores, coloured coded by whether they are contiguous (red) or discontiguous (blue) to EDGE Zone priority cells. The irreplaceability of EDGE Zone cells (priority cells paired with contiguous cells) is shown in panel (b), where irreplaceability refers to the proportional frequency in which cells were selected from 1000 iterations.

**S7 Fig. Threatened evolutionary history prioritisation with complementarity at a resolution of 193 km × 193 km**. Cells were selected iteratively based on those with the highest summed EDGE score using spatial complementarity at a resolution of 193 km 193 km (in red). Priority cells selected using a 96.5 km × 96.5 km resolution are shown in a black outline.

**S8 Fig. Comparison of a branch-length vs a median EDGE score approach for the selection of priority grid cells.** The complementarity procedure was compared across 30 iterations based on the distribution of threat-weighted phylogenetic trees used in Gumbs et al. (2022b); cells were selected based on calculations of expected phylogenetic diversity loss (ePD loss; the branch-length approach) and on species-specific EDGE scores calculated from the same trees. The plot displays the frequency in which cells selected using calculations of ePD loss were also selected using summed EDGE scores, showing a strong correlation using spearman’s rank (ρ = 0.958, p < 0.0001). There was a 97.7% overlap in the location of cells selected between methods.

**S9 Fig. EDGE rarity prioritisation with complementarity.** The map shows priority grid cells based on summed EDGE scores in blue and EDGE rarity scores in red, with sites of mutual overlap shown in black. Here, we define the EDGE rarity of a species as its EDGE score divided by the number of grid cells its range overlaps with. For both metrics, priority cells were selected iteratively with complementarity until the pooled species composition represented 25% of threatened tetrapod evolutionary history.

**S10 Fig. Biodiversity Hotspots.** Biodiversity Hotspots, as defined by Myers et al. (2000), mapped in terms of their (a) threatened evolutionary history (using summed EDGE scores, given in MY) and (b) their EDGE tetrapod species richness at 96.5 km × 96.5 km resolution.

**S11 Fig. Evolutionary Distinctiveness of tetrapod species with absent range data.** Histograms of the Evolutionary Distinctiveness (ED) of species with absent range data visualised across 20 percentiles for (a) amphibians (Median ED = 7.32, Median Percentile = 37th, n = 1215), (b) birds (Median ED = 2.82, Median Percentile = 43rd, n = 35), (c) mammals (Median ED = 1.64, Median Percentile = 40th, n = 522), and (d) reptiles (Median ED = 5.59, Median Percentile = 44th, n = 837).

**S12 Fig. The complementary set of grid cells needed to capture 100% of threatened tetrapod evolutionary history.** 3001 cells, identified iteratively based on the highest scoring summed EDGE values, were selected to capture the threatened evolutionary history of all 33,628 mapped tetrapod species. Cells are coloured coded by the proportion of threatened evolutionary history captured by component cells in 25% intervals.

**S1 Text. A comparison of using summed EDGE scores vs phylogenetic branch lengths for the calculation of threatened evolutionary history.** The (a) overlap in priority sites of threatened evolutionary history calculated using summed EDGE scores vs phylogenetic branch lengths at six different percentiles and (b) the results of a Pearson’s correlation adjusted for spatial autocorrelation.

**S2 Text. Predictors of irreplaceability in EDGE Zone grid cells.** The effects of five variables on the irreplaceability scores of 112 EDGE Zone grid cells, assessed using General Linear Models with a binomial family distribution. Significant *p* values at the 5% criterion are bolded.

**S3 Text. Comparison of the tetrapod threatened evolutionary history within and outside of EDGE Zones.** The (a) median ED scores and (b) median EDGE scores of species from each tetrapod group stratified by whether the grid cell is found within or outside of an EDGE Zone. (c) The proportional contribution each tetrapod group made to the threatened evolutionary history in each grid cell relative to their richness. Here, a higher proportional contribution means a group contributes more to the threatened evolutionary history than expected relative to their richness. Comparisons were made using ANOVA with Tukey’s Honest Significant Difference Test.

## Bibliography

1. Watson JE, Jones KR, Fuller RA, Marco MD, Segan DB, Butchart SH, et al. Persistent disparities between recent rates of habitat conversion and protection and implications for future global conservation targets. Conservation Letters. 2016;9(6):413–21. doi: 10.1111/conl.12295

2. Geldmann J, Manica A, Burgess ND, Coad L, Balmford A. A global-level assessment of the effectiveness of protected areas at resisting anthropogenic pressures. Proceedings of the National Academy of Sciences. 2019;116(46):23209–15. doi: 10.1073/pnas.190822111

3. Maxwell SL, Cazalis V, Dudley N, Hoffmann M, Rodrigues AS, Stolton S, et al. Area-based conservation in the twenty-first century. Nature. 2020;586(7828):217–27. doi: 10.1038/s41586-020-2773-z

4. Myers N, Mittermeier RA, Mittermeier CG, Da Fonseca GA, Kent J. Biodiversity hotspots for conservation priorities. Nature. 2000;403(6772):853. doi: 10.1038/35002501

5. Mittermeier RA, Robles-Gil P, Hoffmann M, Pilgrim J, Brooks T, Mittermeier CG, Lamoreux J, Da Fonseca GAB. Hotspots revisited: Earth’s biologically richest and most endangered terrestrial ecoregions. Mexico City: Cemex; 2004.

6. Díaz S, Zafra-Calvo N, Purvis A, Verburg PH, Obura D, Leadley P, et al. Set ambitious goals for biodiversity and sustainability. Science. 2020;370(6515):411–3. doi: 10.1126/science.abe1530

7. IPBES. Global assessment report on biodiversity and ecosystem services of the Intergovernmental Science-Policy Platform on Biodiversity and Ecosystem Services. Germany: IPBES secretariat; 2019. Available: https://ipbes.net/global-assessment.

8. Gumbs R, Chaudhary A, Daru BH, Faith DP, Forest F, Gray CL, et al. Indicators to monitor the status of the tree of life. Conservation Biology. 2023; Forthcoming. doi: 10.1111/cobi.14141

9. Winter M, Devictor V, Schweiger O. Phylogenetic diversity and nature conservation: where are we? Trends in ecology & evolution. 2013;28(4):199–204. doi: 10.1016/j.tree.2012.10.015

10. Robuchon M, Pavoine S, Véron S, Delli G, Faith DP, Mandrici A, et al. Revisiting species and areas of interest for conserving global mammalian phylogenetic diversity. Nature Communications. 2021;12(1):1–11. doi: 10.1038/s41467-021-23861-y

11. Robuchon M, da Silva J, Dubois G, Gumbs R, Hoban S, Laikre L, et al. Conserving species’ evolutionary potential and history: opportunities under the new post-2020 global biodiversity framework. Authorea Preprints. 2022. doi: 10.22541/au.166568699.98478640/v1

12. Davis AJ, Jenkinson LS, Lawton JH, Shorrocks B, Wood S. Making mistakes when predicting shifts in species range in response to global warming. Nature. 1998;391(6669):783–6. doi: 10.1038/35842

13. Purvis A, Agapow P-M, Gittleman JL, Mace GM. Nonrandom extinction and the loss of evolutionary history. Science. 2000;288(5464):328–30. doi: 10.1126/science.288.5464.328

14. Cadotte MW, Jonathan Davies T. Rarest of the rare: advances in combining evolutionary distinctiveness and scarcity to inform conservation at biogeographical scales. Diversity and Distributions. 2010;16(3):376–85. doi: 10.1111/j.1472-4642.2010.00650.x

15. Forest F, Grenyer R, Rouget M, Davies TJ, Cowling RM, Faith DP, et al. Preserving the evolutionary potential of floras in biodiversity hotspots. Nature. 2007;445(7129):757–60. doi: 10.1038/nature05587

16. Molina-Venegas R, Rodríguez MÁ, Pardo-de-Santayana M, Ronquillo C, Mabberley DJ. Maximum levels of global phylogenetic diversity efficiently capture plant services for humankind. Nature Ecology & Evolution. 2021;5(5):583–8. doi: 10.1038/s41559-021-01414-2

17. Molina-Venegas R. Conserving evolutionarily distinct species is critical to safeguard human well-being. Scientific reports. 2021;11(1):1–9. doi: 10.1038/s41598-021-03616-x

18. Faith DP. Conservation evaluation and phylogenetic diversity. Biological conservation. 1992;61(1):1–10. doi: 10.1016/0006-3207(92)91201-3

19. Faith DP, Veron S, Pavoine S, Pellens R. Indicators for the Expected Loss of Phylogenetic Diversity. In: Scherson RA, Faith DP, editors. Phylogenetic Diversity: Applications and Challenges in Biodiversity Science. Cham: Springer International Publishing; 2018. pp. 73–91. doi: 10.1007/978-3-319-93145-6_4

20. Owen NR, Gumbs R, Gray CL, Faith DP. Global conservation of phylogenetic diversity captures more than just functional diversity. Nature communications. 2019;10(1):1–3. doi: 10.1038/s41467-019-08600-8

21. Fritz SA, Rahbek C. Global patterns of amphibian phylogenetic diversity. Journal of biogeography. 2012;39(8):1373–82. doi: 10.1111/j.1365-2699.2012.02757.x

22. Safi K, Cianciaruso MV, Loyola RD, Brito D, Armour-Marshall K, Diniz-Filho JAF. Understanding global patterns of mammalian functional and phylogenetic diversity. Philosophical Transactions of the Royal Society B: Biological Sciences. 2011;366(1577):2536–44. doi: 10.1098/rstb.2011.0024

23. Safi K, Armour-Marshall K, Baillie JE, Isaac NJ. Global patterns of evolutionary distinct and globally endangered amphibians and mammals. PloS one. 2013;8(5):e63582. doi: 10.1371/journal.pone.0063582

24. Daru BH, van der Bank M, Davies TJ. Spatial incongruence among hotspots and complementary areas of tree diversity in southern A frica. Diversity and Distributions. 2015;21(7):769–80. doi: 10.1111/ddi.12290

25. Pollock LJ, Thuiller W, Jetz W. Large conservation gains possible for global biodiversity facets. Nature. 2017;546(7656):141–4. doi: 10.1038/nature22368

26. Brum FT, Graham CH, Costa GC, Hedges SB, Penone C, Radeloff VC, et al. Global priorities for conservation across multiple dimensions of mammalian diversity. Proceedings of the National Academy of Sciences. 2017;114(29):7641–6. doi: 10.1073/pnas.1706461114

27. Rosauer D, Laffan SW, Crisp MD, Donnellan SC, Cook LG. Phylogenetic endemism: a new approach for identifying geographical concentrations of evolutionary history. Molecular ecology. 2009;18(19):4061–72. doi: 10.1111/j.1365-294X.2009.04311.x

28. Murali G, Gumbs R, Meiri S, Roll U. Global determinants and conservation of evolutionary and geographic rarity in land vertebrates. Science advances. 2021;7(42):eabe5582. doi: 10.1126/sciadv.abe5582

29. Gumbs R, Gray CL, Böhm M, Hoffmann M, Grenyer R, Jetz W, et al. Global priorities for conservation of reptilian phylogenetic diversity in the face of human impacts. Nature communications. 2020;11(1):1–13. doi: 10.1038/s41467-020-16410-6

30. Daru BH, le Roux PC, Gopalraj J, Park DS, Holt BG, Greve M. Spatial overlaps between the global protected areas network and terrestrial hotspots of evolutionary diversity. Global Ecology and Biogeography. 2019;28(6):757–66. doi: 10.1111/geb.12888

31. Rapacciuolo G, Graham CH, Marin J, Behm JE, Costa GC, Hedges SB, et al. Species diversity as a surrogate for conservation of phylogenetic and functional diversity in terrestrial vertebrates across the Americas. Nature ecology & evolution. 2019;3(1):53–61. doi: 10.1038/s41559-018-0744-7

32. Sechrest W, Brooks TM, da Fonseca GA, Konstant WR, Mittermeier RA, Purvis A, et al. Hotspots and the conservation of evolutionary history. Proceedings of the National Academy of Sciences. 2002;99(4):2067–71. doi: 10.1073/pnas.251680798

33. Isaac NJ, Turvey ST, Collen B, Waterman C, Baillie JE. Mammals on the EDGE: conservation priorities based on threat and phylogeny. PloS one. 2007;2(3):e296. doi: 10.1371/journal.pone.0000296

34. Collen B, Turvey ST, Waterman C, Meredith HM, Kuhn TS, Baillie JE, et al. Investing in evolutionary history: implementing a phylogenetic approach for mammal conservation. Philosophical Transactions of the Royal Society B: Biological Sciences. 2011;366(1578):2611–22. doi: 10.1098/rstb.2011.0109

35. Isaac NJ, Redding DW, Meredith HM, Safi K. Phylogenetically-informed priorities for amphibian conservation. 2012. doi: 10.1371/journal.pone.0043912

36. Jetz W, Thomas GH, Joy JB, Redding DW, Hartmann K, Mooers AO. Global distribution and conservation of evolutionary distinctness in birds. Current biology. 2014;24(9):919–30. doi: 10.1016/j.cub.2014.03.011

37. Curnick D, Head C, Huang D, Crabbe MJC, Gollock M, Hoeksema B, et al. Setting evolutionary-based conservation priorities for a phylogenetically data-poor taxonomic group (S cleractinia). Animal Conservation. 2015;18(4):303–12. doi: 10.1111/acv.12185

38. Gumbs R, Gray CL, Wearn OR, Owen NR. Tetrapods on the EDGE: Overcoming data limitations to identify phylogenetic conservation priorities. PLoS One. 2018;13(4):e0194680. doi: 10.1371/journal.pone.0194680

39. Forest F, Moat J, Baloch E, Brummitt NA, Bachman SP, Ickert-Bond S, et al. Gymnosperms on the EDGE. Scientific reports. 2018;8(1):1–11. doi: 10.1038/s41598-018-24365-4

40. Stein RW, Mull CG, Kuhn TS, Aschliman NC, Davidson LN, Joy JB, et al. Global priorities for conserving the evolutionary history of sharks, rays and chimaeras. Nature ecology & evolution. 2018;2(2):288–98. doi: 10.1038/s41559-017-0448-4

41. Gumbs R, Gray CL, Bohm M, Burfield IJ, Couchman OR, Faith DP, et al. The EDGE2 protocol: Advancing the prioritisation of Evolutionarily Distinct and Globally Endangered species for practical conservation action. PLOS Biology. 2023;21(2):e3001991. doi: 10.1371/journal.pbio.3001991

42. Steel M, Mimoto A, Mooers AØ. Hedging our bets: the expected contribution of species to future phylogenetic diversity. Evolutionary Bioinformatics. 2007;3:117693430700300024. doi: 10.1177/117693430700300024

43. Faith DP. Threatened species and the potential loss of phylogenetic diversity: conservation scenarios based on estimated extinction probabilities and phylogenetic risk analysis. Conservation Biology. 2008;22(6):1461–70. doi: 10.1111/j.1523-1739.2008.01068.x

44. May-Collado LJ, Zambrana-Torrelio C, Agnarsson I. Global spatial analyses of phylogenetic conservation priorities for aquatic mammals. In: Pellens R, Grandcolas P. Biodiversity conservation and phylogenetic systematics: Springer, Cham; 2016. p. 305–18. doi: 10.1007/978-3-319-22461-9_15

45. Mishler BD, Knerr N, González-Orozco CE, Thornhill AH, Laffan SW, Miller JT. Phylogenetic measures of biodiversity and neo-and paleo-endemism in Australian Acacia. Nature Communications. 2014;5(1):1–10. doi: 10.1038/ncomms5473

46. Chaudhary A, Pourfaraj V, Mooers AO. Projecting global land use-driven evolutionary history loss. Diversity and Distributions. 2018;24(2):158–67. doi: 10.1111/ddi.12677

47. Roll U, Feldman A, Novosolov M, Allison A, Bauer AM, Bernard R, et al. The global distribution of tetrapods reveals a need for targeted reptile conservation. Nature ecology & evolution. 2017;1(11):1677–82. doi: 10.1038/s41559-017-0332-2

48. Vallejos R, Osorio F, Bevilacqua M. Spatial relationships between two georeferenced variables: With applications in R: Springer Nature; 2020. doi: 10.1007/978-3-030-56681-4

49. Dutilleul P, Clifford P, Richardson S, Hemon D. Modifying the t test for assessing the correlation between two spatial processes. Biometrics. 1993:305–14. doi: 10.2307/2532625

50. Redding DW, Mooers AØ. Incorporating evolutionary measures into conservation prioritization. Conservation Biology. 2006;20(6):1670–8. doi: 10.1111/j.1523-1739.2006.00555.x

51. Cadotte MW, Tucker CM. Difficult decisions: Strategies for conservation prioritization when taxonomic, phylogenetic and functional diversity are not spatially congruent. Biological conservation. 2018;225:128–33. doi: 10.1016/j.biocon.2018.06.014

52. Alkire S, Kanagaratnam U, and Suppa N. The global Multidimensional Poverty Index (MPI) 2021’, OPHI MPI Methodological Note 51. Oxford Poverty and Human Development Initiative, University of Oxford. 2021.

53. Venter O, Sanderson EW, Magrach A, Allan JR, Beher J, Jones KR, et al. Sixteen years of change in the global terrestrial human footprint and implications for biodiversity conservation. Nature communications. 2016;7(1):1–11. doi: 10.1038/ncomms12558

54. Convention on Biological Diversity. Strategic Plan for Biodiversity 2011-2020, including Aichi Biodiversity Targets. 2011. Available: https://www.cbd.int/doc/strategic-plan/2011-2020/Aichi-Targets-EN.pdf

55. Conference of the Parties to the Convention on Biological Diversity. Decision adopted by the Conference of the Parties to the Convention on Biological Diversity. In: CBD/COP/DEC/15/4 [Internet]. 2022. Available: https://www.cbd.int/doc/decisions/cop-15/cop-15-dec-04-en.pdf

56. Ganzhorn JU, Lowry PP, Schatz GE, Sommer S. The biodiversity of Madagascar: one of the world’s hottest hotspots on its way out. Oryx. 2001;35(4):346–8. doi: 10.1046/j.1365-3008.2001.00201.x

57. Harfoot MB, Johnston A, Balmford A, Burgess ND, Butchart SH, Dias MP, et al. Using the IUCN Red List to map threats to terrestrial vertebrates at global scale. Nature Ecology & Evolution. 2021;5(11):1510–9. doi: 10.1038/s41559-021-01542-9

58. Vences M, Wollenberg KC, Vieites DR, Lees DC. Madagascar as a model region of species diversification. Trends in ecology & evolution. 2009;24(8):456–65. doi: 10.1016/j.tree.2009.03.011

59. Wilmé L, Goodman SM, Ganzhorn JU. Biogeographic evolution of Madagascar’s microendemic biota. science. 2006;312(5776):1063–5. doi: 10.1126/science.1122806

60. Sodhi NS, Koh LP, Brook BW, Ng PK. Southeast Asian biodiversity: an impending disaster. Trends in ecology & evolution. 2004;19(12):654–60. doi: 10.1016/j.tree.2004.09.006

61. Tilman D, Clark M, Williams DR, Kimmel K, Polasky S, Packer C. Future threats to biodiversity and pathways to their prevention. Nature. 2017;546(7656):73–81. doi: 10.1038/nature22900

62. Tchassem F AM, Doherty-Bone TM, Kameni N MM, Tapondjou N WP, Tamesse JL, Gonwouo LN. What is driving declines of montane endemic amphibians? New insights from Mount Bamboutos, Cameroon. Oryx. 2021;55(1):23–33. doi: 10.1017/S0030605318001448

63. Doherty-Bone TM, Gvoždík V. The Amphibians of Mount Oku, Cameroon: an updated species inventory and conservation review. ZooKeys. 2017 (643):109. doi: 10.3897/zookeys.643.9422

64. Stuart SN, Chanson JS, Cox NA, Young BE, Rodrigues AS, Fischman DL, et al. Status and trends of amphibian declines and extinctions worldwide. Science. 2004;306(5702):1783–6. doi: 10.1126/science.1103538

65. Hedges SB, Díaz LM. The conservation status of amphibians in the West Indies. In: Hailey A, Wilson BS, Horrocks JA. Conservation of Caribbean Island Herpetofaunas Volume 1: Conservation Biology and the Wider Caribbean: Brill; 2011. p. 31–47. doi: 10.1163/ej.9789004183957.i-228

66. Mittermeier RA, Turner WR, Larsen FW, Brooks TM, Gascon C. Global biodiversity conservation: the critical role of hotspots. Biodiversity Hotspots: Springer; 2011. p. 3–22. doi: 10.1007/978-3-642-20992-5_1

67. Sloan S, Jenkins CN, Joppa LN, Gaveau DL, Laurance WF. Remaining natural vegetation in the global Biodiversity Hotspots. Biological Conservation. 2014;177:12–24. doi: 10.1016/j.biocon.2014.05.027

68. Habel JC, Rasche L, Schneider UA, Engler JO, Schmid E, Rödder D, et al. Final countdown for Biodiversity Hotspots. Conservation Letters. 2019;12(6):e12668. doi: 10.1111/conl.12668

69. Myers N. Threatened biotas: “hot spots” in tropical forests. Environmentalist. 1988;8(3):187–208. doi: 10.1007/BF02240252

70. Shearman PL, Ash J, Mackey B, Bryan JE, Lokes B. Forest conversion and degradation in Papua New Guinea 1972–2002. Biotropica. 2009;41(3):379–90. doi: 10.1111/j.1744-7429.2009.00495.x

71. Wyborn C, Evans MC. Conservation needs to break free from global priority mapping. Nature Ecology & Evolution. 2021;5(10):1322–4. doi: 10.1038/s41559-021-01540-x

72. Baker WJ, Bailey P, Barber V, Barker A, Bellot S, Bishop D, et al. A comprehensive phylogenomic platform for exploring the angiosperm tree of life. Systematic biology. 2022;71(2):301–19. doi: 10.1093/sysbio/syab035

73. Nic Lughadha E, Bachman SP, Leão TC, Forest F, Halley JM, Moat J, et al. Extinction risk and threats to plants and fungi. Plants, People, Planet. 2020;2(5):389–408. doi: 10.1002/ppp3.10146

74. Gumbs R, Scott O, Bates R, Böhm M, Forest F, Gray C, et al. Global conservation status of the jawed vertebrate Tree of Life. Research Square [Preprint]. 2023 [cited 2023 July 18]. Available from: https://www.researchsquare.com/article/rs-2835356/v1 doi: 10.21203/rs.3.rs-2835356/v1

75. IUCN. IUCN Red List of Threatened Species. Version 2020-1. 2020. Available: www.iucnredlist.org

76. BirdLife International and Handbook of the Birds of the World. Bird species distribution maps of the world. Version 2020.1. 2020. Available: http://datazone.birdlife.org/species/requestdis

77. ZSL EDGE of Existence. EDGE of Existence. 2022. Available: http://edgeofexistence.org/

78. Noss RF, Platt WJ, Sorrie BA, Weakley AS, Means DB, Costanza J, et al. How global biodiversity hotspots may go unrecognized: lessons from the North American Coastal Plain. Diversity and Distributions. 2015;21(2):236–44. doi: 10.1111/ddi.12278

79. Olson DM, Dinerstein E, Wikramanayake ED, Burgess ND, Powell GV, Underwood EC, et al. Terrestrial Ecoregions of the World: A New Map of Life on Earth. BioScience. 2001;51(11):933–8. doi: 10.1641/0006-3568(2001)051[0933:TEOTWA]2.0.CO;2

80. Williams BA, Venter O, Allan JR, Atkinson SC, Rehbein JA, Ward M, et al. Change in terrestrial human footprint drives continued loss of intact ecosystems. One Earth. 2020;3(3):371–82. doi: 10.1016/j.oneear.2020.08.009

81. Di Marco M, Venter O, Possingham HP, Watson JE. Changes in human footprint drive changes in species extinction risk. Nature communications. 2018;9(1):1–9. doi: 10.1038/s41467-018-07049-5

82. Allan JR, Venter O, Maxwell S, Bertzky B, Jones K, Shi Y, et al. Recent increases in human pressure and forest loss threaten many Natural World Heritage Sites. Biological conservation. 2017;206:47–55. doi: 10.1016/j.biocon.2016.12.011

83. IUCN, UNEP-WCMC. The World Database on Protected Areas (WDPA). UNEP World Conservation Monitoring Centre, Cambridge (UK): Version 1.6: 2021 [accessed June 2021]. Available: www.protectedplanet.net

